# Temporal control of cortico-thalamic neuron specification by regulation of Neurogenin activity and Polycomb repressive complexes

**DOI:** 10.1101/431684

**Authors:** Koji Oishi, Debbie L. C. van den Berg, Franç Guillemot

## Abstract

Neural progenitor cells (NPCs) in the embryonic mammalian neocortex generate different neuronal subtypes sequentially. A long-standing hypothesis to account for this temporal fate specification process is that NPCs change their differentiation potential over time. However, the molecular mechanisms underlying these temporal changes in NPC properties are poorly understood. Here we show that Neurogenin1 and Neurogenin2 (Neurog1/2), two proneural transcription factors expressed in NPCs throughout cortical neurogenesis, specify the identity of one of the first cortical neuron subtypes generated, layer 6 cortico-thalamic neurons (CTNs). We found that *Neurog1/2* specify the CTN fate through regulation of the cortical fate determinants *Fezf2* and *Foxp2* and that this *Neurog*-induced programme becomes inactive after the period of CTN production. Two independent mechanisms contribute to the arrest of CTN neuron generation at the end of layer 6 neurogenesis, including a reduction in the transcriptional activity of Neurog1/2 and the deposition of epigenetic repressive modifications mediated by Polycomb repressive complexes at the *Foxp2* gene. Therefore, the duration of production of a cortical neuron subtype is controlled by multiple locking mechanisms involving both transcriptional and epigenetic processes.

## Introduction

Neural stem cells and intermediate progenitors (called thereafter neural progenitor cells or NPCs) which reside in the ventricular and subventricular zones (VZ/SVZ) of the mammalian neocortex, give rise to distinct subtypes of neurons with specific cell morphologies, birth dates and connections with other regions of the nervous system (Molyneaux et al., 2007; Lodato and Arlotta, 2015; Martynoga et al., 2012). Cortical neurons are born either directly from stem cells in the VZ or indirectly from intermediate progenitors which reside in the cortical SVZ (Florio and Huttner, 2014; Sun and Hevner, 2014). They are generated in a sequential manner, with the earliest NPCs generating subplate neurons around E11.5 while subsequent NPCs produce neurons in layers 6, 5, 4, and eventually 2/3 around E12.5, E13.5, E14.5 and E15.5, respectively. This crucial feature of cortical development has led to a model whereby cortical neurons acquire their distinct laminar identities by temporal specification of cortical NPCs, with the differentiation potential of NPCs changing over time (Greig et al., 2013; McConnell, 1995), akin to the temporal specification of neuroblasts in the ventral nerve cord and optic lobe of Drosophila (Kohwi and Doe, 2013). However, there is scant evidence that NPCs are temporally specified during mammalian corticogenesis and few temporally-controlled factors that might drive such a process have been identified (Alsio et al., 2013; Dominguez et al., 2013; Frantz et al., 1994; Hirata et al., 2004; Zahr et al., 2018).

Studies from Sue McConnell’s group involving heterochronic transplantation experiments have demonstrated that cortical NPCs undergo a progressive restriction of fate potential during cortical development. They showed that early NPCs, producing layer 6 neurons, could differentiate into upper layer neurons when transplanted in the late VZ, while late NPCs, producing layer 2/3 neurons did not differentiate into deep layer neurons when transplanted into the early VZ (Frantz and McConnell, 1996; McConnell and Kaznowski, 1991). Recent studies have identified molecular determinants that are critical for the specification of particular cortical neuron subtypes, but most of these factors are expressed only in postmitotic neurons [e.g. Bcl11b (Arlotta et al., 2005); Bhlhb5 (Joshi et al., 2008); Rorb (Oishi et al., 2016a); Pcdh20 (Oishi et al., 2016b); Zfpm2 (Galazo et al., 2016)] or in both NPCs and neurons [Fezf2 (Hirata et al., 2004; Molyneaux et al., 2005), Pou3fs (Dominguez et al., 2013), Otx1 (Frantz et al., 1994)]. It remains therefore unclear whether cell fate determination actually occurs in NPCs as posited by the temporal specification hypothesis (Alsio et al., 2013).

We previously reported that Neurog1/2, two proneural basic helix-loop-helix (bHLH) transcription factors that are expressed in NPCs and only transiently in postmitotic neurons, promote the generation and neuronal differentiation of cortical neurons but also the acquisition of aspects of the subtype identity of deep layer (early born) neurons, and not that of upper layer (late born) neurons (Schuurmans et al., 2004). However, the downstream mechanisms through which *Neurog1/2* specify a deep layer fate have not yet been determined. Moreover, the fact that *Neurog1/2* are prominently expressed by NPCs throughout the neurogenic period although they specify neuronal subtypes only during the early stages of neurogenesis, raises the question of how this fate specification mechanism is temporally regulated.

Epigenetic control of gene expression, and particularly chromatin remodeling by modifications of histones, have been shown to play pivotal roles in cell fate specification in a wide variety of cell types including NPCs (Grossniklaus and Paro, 2014; Margueron and Reinberg, 2011; Yoon et al., 2018). Polycomb repressive complexes (PRCs) in particular have been implicated in the regulation of cerebral cortex development, including in the timing of generation of particular cortical cell types. PRCs are composed of PRC2, which deposits methyl residues on Lys27 in Histone H3 (H3K27me3) and PRC1, which recognizes H3K27me3 and ubiquitinates Lys119 in Histone H2A (H2AK119ub) (Blackledge et al., 2015; Simon and Kingston, 2013). Ezh2, a catalytic component of PRC2, has been reported to play a role in the balance between self-renewal and differentiation of cortical NPCs (Pereira et al., 2010). Moreover, in the absence of *Ezh2,* many subtype-specific markers are expressed prematurely in the early cortex (Pereira et al., 2010), suggesting a global repression of subtype specific genes by PRCs. More recently, Ring1B, a catalytic subunit of PRC1, has been implicated in the restriction of the differentiation potential of cortical NPCs. *Ring1B* knockout was reported to extend the period of generation of layer 5 neurons and to reduce the period of generation of upper layer neurons, suggesting that PRC1 is required for the termination of layer 5 production (Morimoto-Suzki et al., 2014). How widespread is the role of PRCs in the temporal specification of cortical neuron subtypes is however still unclear and whether they control the timing of *Neurog1/2* activity of deep layer neuron specification is unknown.

In this study, we have further defined the role of *Neurog1/2* in specification of deep layer cortical neurons and we have identified the mechanisms that restrict temporally this activity without affecting *Neurog1/2* proneural function that operates throughout cortical development. We found that changes in both Neurog1/2 transcriptional activity and PRC-dependent regulation of some of their targets restrict *Neurog1/2* subtype specification function during cortical development.

## Results

### Failure of generation of cortico-thalamic neurons in *Neurogenin* mutant mice

Our previous work showed that *Neurog1/2* are required for the correct specification of several aspects of deep layer cortical neuron identities but not for the specification of superficial layer neuronal identities (Schuurmans et al., 2004). However, because of a lack of appropriate markers at the time, it was unclear which particular subtypes of deep layer neurons, which include layer 5 subcerebral (e.g. cortico-spinal, cortico-pontine) neurons, layer 6 cortico-thalamic neurons (CTNs) and layer 6 and layer 5 cortico-cortical (CCNs) neurons, require *Neurog1/2* for their specification. To better define the role of *Neurog1/2* in cortical neuron subtype specification, we re-examined the neocortex of *Neurog1/2* mutant mice using markers specific for each of the different subtypes of deep layer neurons.

Bcl11b (also known as Ctip2), a well-established marker for subcortical neurons, is expressed strongly by layer 5 subcerebral neurons and at a lower level by layer 6 CTNs (Arlotta et al., 2005). We observed a decrease (46% decrease) in the number of Bcl11b-positive cells in layer 6, corresponding to CTNs, but not in layer 5, corresponding to subcerebral neurons, in *Neurog2* mutant mice and more strongly in *Neurog1/2* mutants (64% decrease) at E18.5 (Figures 1A–1C and 1M). Moreover, the number of Satb2-positive cells in layer 5, corresponding to layer 5 CCNs, remained unchanged in *Neurog2* and *Neurog1/2* mutants (Figures 1A–1C and 1N). This finding prompted us to investigate in detail the specification of layer 6 neurons in these mice. We first examined whether the entire layer 6 neuronal population was affected in *Neurog1/2* mutants. The number of cells in layer 6, visualized by nuclear staining, was slightly reduced in *Neurog2* and *Neurog1/2* mutants (16% and 17% decrease, respectively) (Figure 1O). However, the extent of reduction of Bcl11b-positive cells in layer 6 was more profound than that of all layer 6 neurons (46% versus 16% decrease), and the percentage of Bcl11b-positive cells among layer 6 neurons was decreased in *Neurog2* and *Neurog1/2* mutants (36% and 56% decrease, respectively) (Figure 1P).

**Figure 1.**
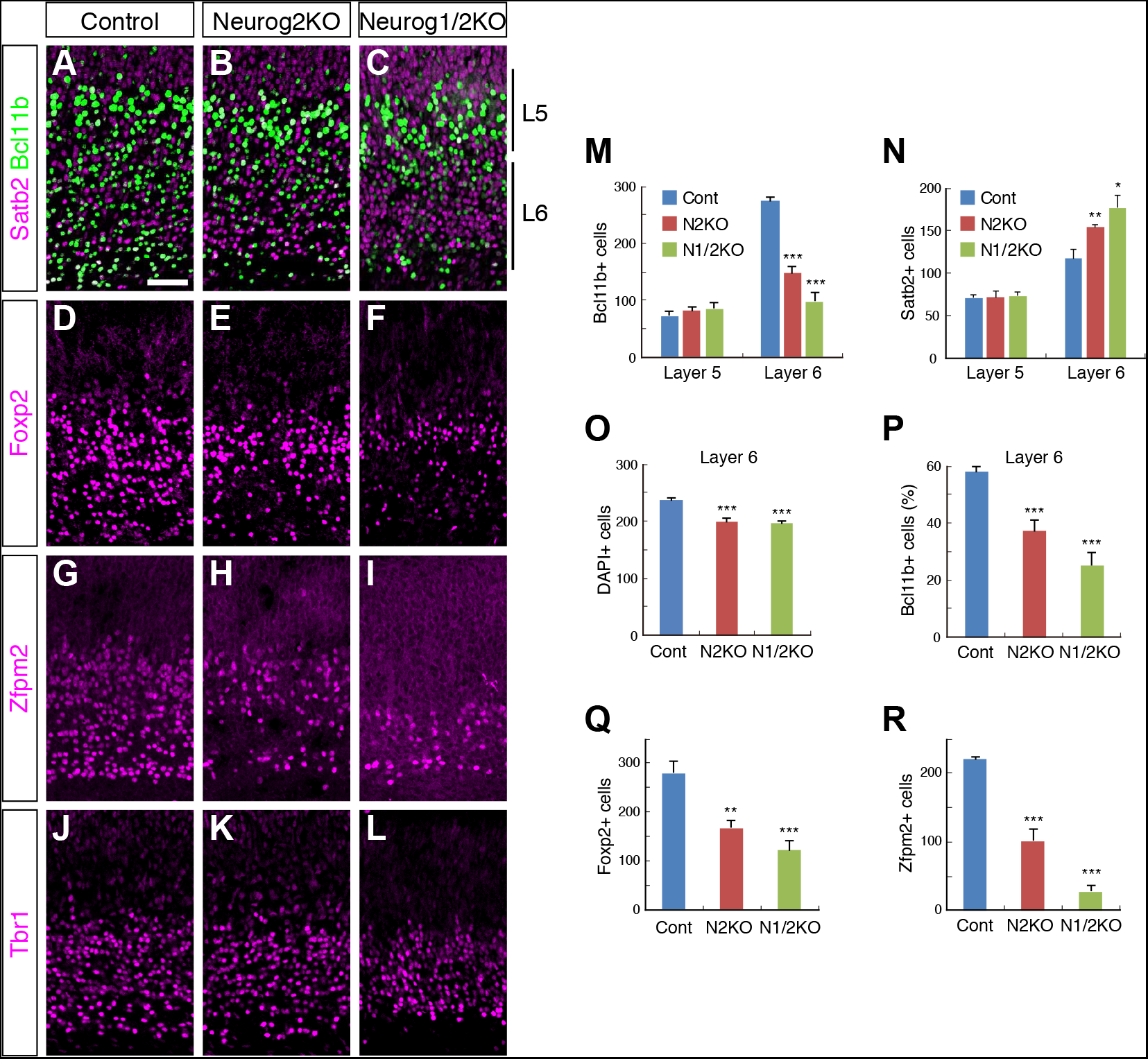
Inactivation of *Neurog1/2* decreases generation of cortico-thalamic neurons. (A–L) Immunostaining for the indicated subtype-specific makers in E18.5 cortices of control, *Neurog2KO,* and *Neurog1/2KO* mice. Tbr1 is generally used as a marker for CTNs but we observed that Tbr1 is also expressed by CCNs at moderate levels (data not shown). (M and N) The numbers of Bcl11b (M)- or Satb2 (N)-positive cells in 220-μm width cortical columns of the indicated genotypes was counted (Control, n = 4; *Neurog2KO,* n = 4; *Neurog1/2KO,* n = 3). (O) The number of cells visualized with DAPI in layer 6 of the indicated genotypes was counted (Control, n = 4; *Neurog2KO,* n = 4; *Neurog1/2KO,* n = 3). (P) Quantification of the percentage of Bcl11b-postive cells in layer 6 (Control, n = 4; *Neurog2KO,* n = 4; *Neurog1/2KO,* n = 3). (Q and R) The number of Foxp2 (Q)- or Zfpm2 (R)-positive cells in the indicated genotypes was counted (Control, n = 4; *Neurog2KO,* n = 4; *Neurog1/2KO,* n = 4). Scale bar: 50 μm. Data are represented as means ± SD; \**p* < 0.005, \*\**p* < 0.001, \*\*\**p* < 0.0001 from two-tailed unpaired Student’s t-test with Welch’s correction.

As Bcl11b was reported to label CTNs, we also examined the expression of other CTN makers, including the transcription factors Foxp2 (Hisaoka et al., 2010), Zfpm2 (Han et al., 2011), Nfia (Betancourt et al., 2014) and the signal transduction protein Ppp1r1b/Darpp32 (Molyneaux et al., 2005). The numbers of cells expressing these markers were all reduced in *Neurog2* and *Neurog1/2* mutant mice (Figures 1D–1I, 1Q and 1R; Figures S1A–S1F and S1M), confirming that *Neurog1* and *Neurog2* are required for the generation of CTN neurons.

### Fate switch of *Neurogenin-*deficient cortico-thalamic neurons to cortico-cortical neurons

The observed reduction in percentage of CTNs among layer 6 neurons in *Neurog* mutant mice suggested that some CTNs might be mis-specified and acquire alternative identities in the absence of *Neurog1/2.* CCNs, which send axons to the contralateral or ipsilateral cortex, constitute another major subtype of layer 6 neurons (Thomson, 2010; Tasic et al., 2016), which can be distinguished from CTNs by the expression of Satb2 and Nurr1 (Arimatsu et al., 2003; Watakabe et al., 2007). The numbers of Satb2- and Nurr1-positive cells in layer 6 was indeed increased in E18.5 *Neurog2* and *Neurong1/2* mutant mice (Satb2, 31% and 51%, respectively; Nurr1, 46% and 77%, respectively) (Figures 1A–1C and 1N; Figures S1G–S1I and S1N). Importantly, all cells located in layer 6 of *Neurog* mutant cortices retained molecular characteristics of layer 6 neurons, since they expressed Tbr1, a pan-layer 6 marker (Figures 1J–1L; (Galazo et al., 2016)) and Sox5, a maker of all layer 6 and a subset of layer 5 neurons (Figures S1J–S1L). Therefore, a large fraction of layer 6 neurons fail to be specified into CTNs and switch to another layer 6 subtype, CCNs, in the absence of *Neurog1/2* function.

### *Neurog2* overexpression promotes cortico-thalamic neuron specification

We next asked whether *Neurog1/2* expression was not only required but also sufficient to induce a CTN fate in NPCs destined to produce layer 6 neurons. To address this question, we used *in utero* electroporation (IUE) to express *Neurog2* and the reporter *EGFP* in cortical VZ cells at around E12.5, when most layer 6 neurons are born (Figure 2A). In a control experiment where EGFP alone was electroporated, 82.2 ± 1.8% of EGFP labeled cells analysed at E18.5 were located in layer 6 and 40–50% of the electroporated cells co-expressed CTN markers including Foxp2 (Figures 2A and 2G), Zfpm2 (Figure 2C and 2H), Bcl11b (Figures S2A and S2E) and Nfia (Figures S2C and S2F), while 48.6 ± 1.8% expressed the CCN marker Satb2 (Figures 2E and 2I). Co-expression of *Neurog2* with *EGFP* markedly increased the percentages of CTN marker-positive cells to 75 – 90% (Figures 2B, 2D, 2G, and 2H; Figures S2B and S2D–S2F), while decreasing the percentage of Satb2-positive CCNs to 14.4 ± 3.4% (Figures 2F and 2I). Importantly, the axons of neurons electroporated with *Neurog2* + *EGFP* were more abundant in the dorsal thalamus, corresponding to the trajectory of CTN axons, than those from control neurons (Figures 2J–2N), suggesting that *Neurog2* overexpression changed not only marker gene expression but also the axonal projection pattern of electroporated neurons to a CTN-like phenotype.

**Figure 2.**
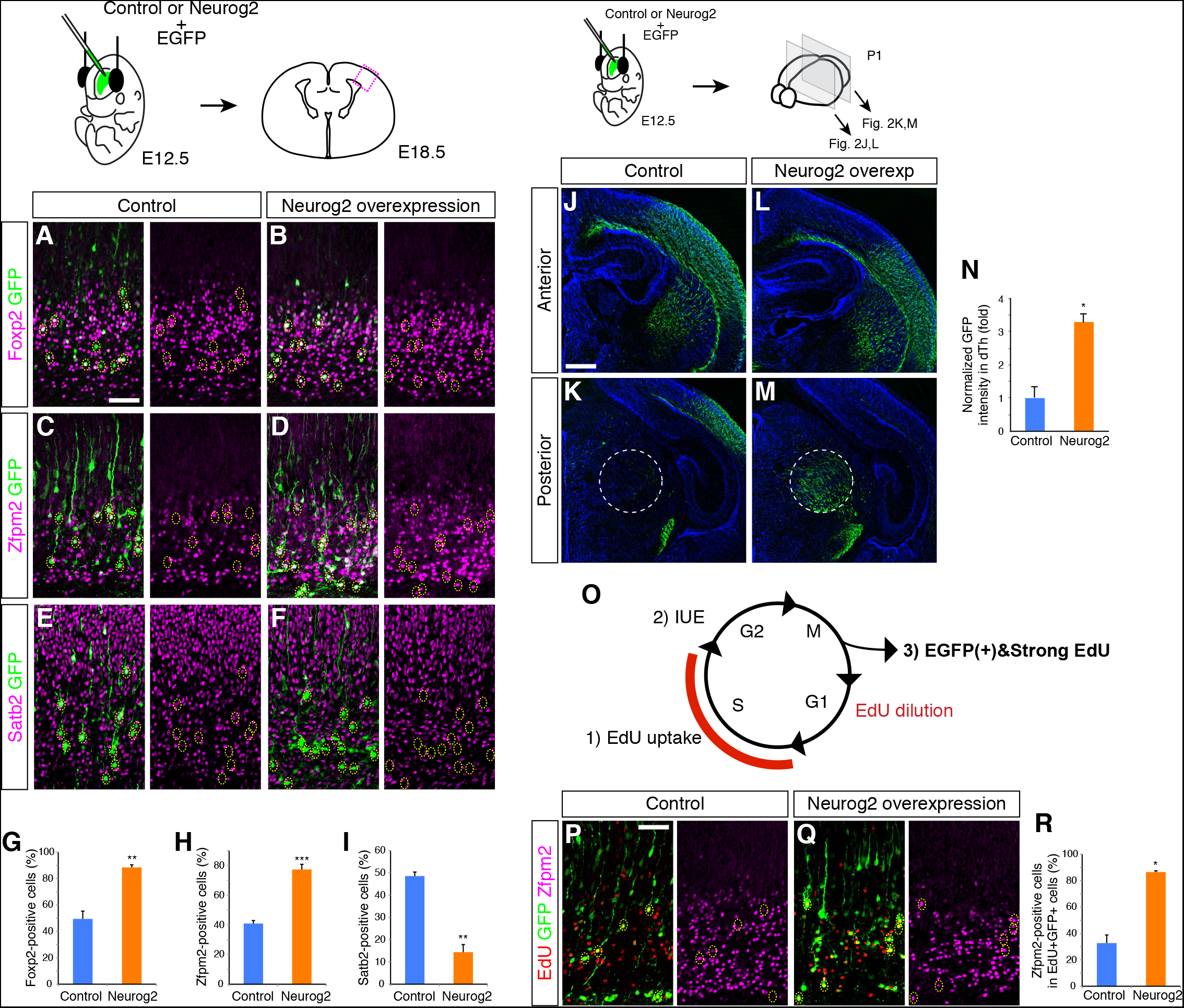
*Neurog2* overexpression promotes cortico-thalamic neuron specification. (A–F) Immunostaining for the indicated subtype-specific markers in E18.5 cortices after *in utero* electroporation at E12.5 of *EGFP* and control (A, C, and E) or *Neurog2* expression vector (B, D, and F). (G–I) Quantification of the percentage of marker-positive cells among EGFP-positive cells in (A)–(F) (Control, n = 4; *Neurog2,* n = 4). (J–M) Immunostaining for EGFP in P1 brains after *in utero* electroporation as in (A) and (B). A representative image is shown in each condition. An anterior level of the electroporated brains is presented to demonstrate electroporation efficiency (J and L). A posterior level including the dorsal thalamus (indicated with dotted circles) is presented to show the axon trajectories of the electroporated cells (K and M). Quantification of (J–M) is shown (N) (Control, n = 4; *Neurog2,* n = 3). (O–R) EdU was administrated 1 hour before *in utero* electroporation at E12.5, and the brains were analyzed at E18.5 (O). EdU staining together with EGFP immunostaining (left panels) and Zfpm2 immunostaining (right panels) are shown (P and Q). The percentage of Zfpm2-positive cells among both EdU- and EGFP-positive cells shown in (P) and (Q) was calculated (R) (Control, n = 4; *Neurog2,* n = 3). Scale bars: 50 μm (A and P); 500 μm (J). Data are represented as means ± SEM; \**p* < 0.005; \*\**p* < 0.001; \*\*\**p* < 0.0001 from two-tailed unpaired Student’s *t*-test with Welch’s correction.

*Neurog* overexpression has been shown to promote cell cycle exit and neuronal differentiation (Hand et al., 2005; Nieto et al., 2001). The *Neurog2*-induced conversion of CCNs into CTNs could in principle be due to the premature differentiation of *Neurog2-* electroporated neurons if CTNs were normally generated before CCNs. To exclude this possibility, we compared Neurog2-electroporated and control *EGFP*-electroporated neurons that were born at the same time by administering EdU one hour before IUE and focusing our analysis on the population of EGFP+ cells that were strongly labeled with EdU, i.e. cells that were NPCs in S phase of the cell cycle at the time of IUE and became postmitotic immediately thereafter (Figure 2O). *Neurog2* overexpression in this population markedly increased the percentage of Zfpm2-positive CTNs compared with the percentage of Zfpm2-expressing CTNs born at the same time in the control electroporation experiment (Figures 2P–2R), suggesting that *Neurog2* directly promotes the CTN fate rather than controlling the timing of layer 6 neuron differentiation. Therefore, expression of *Neurogenins* during the generation of layer 6 neurons is sufficient to promote CTN specification at the expense of CCN specification.

### Identification of target genes of *Neurog2* involved in cortico-thalamic neuron specification

*Neurogenin* genes specify CTNs presumably by inducing CTN-specific target genes in NPCs. To identify such targets, we mined multiple datasets, including microarray data for genes expressed in *Neurog2* mutant, *Neurog1/2* mutant and control E13.5 cortices (Gohlke et al., 2008), ChIP-Seq data for genomic locations bound by Neurog2 in E14.5 cortex (Sessa et al., 2016), publicly available *in situ* hybridization resources including the Allen Developing Mouse Brain Atlas (http://developingmouse.brain-map.org) and BGEM (Magdaleno et al., 2006), and literature searches for genes expressed in mouse cortical layer 6 neurons. Since *Neurog1/2* are expressed in NPCs, we speculated that candidate targets that can execute CTN specification in neurons should include transcription factors that are expressed in NPCs and/or postmitotic neurons. Using these datasets, we identified several relevant candidate *Neurog* target genes, i.e. genes encoding transcription factors that are down-regulated in *Neurog2* and *Neurog1/2* mutant cortices and have Neurog2 ChIP-Seq peaks in their vicinity (Figure 3A). We narrowed further the candidate targets list by examining the effect of *Neurog2* overexpression, which is expected to increase the expression of *bona fide* targets since *Neurog2* has been shown to act as a transcriptional activator (Castro et al., 2006). EGFP-positive cells from cortices electroporated with *Neurog2* plus *EGFP* or *EGFP* alone at E12.5 and dissociated at E16.5 were purified by FACS and the mRNA was prepared for qRT-PCR analysis. RNA levels for some of the candidate targets including *Bcl11b, Fezf2, Nfia* and *Foxp2* were significantly up-regulated by *Neurog2* overexpression (Figure 3B).

**Figure 3.**
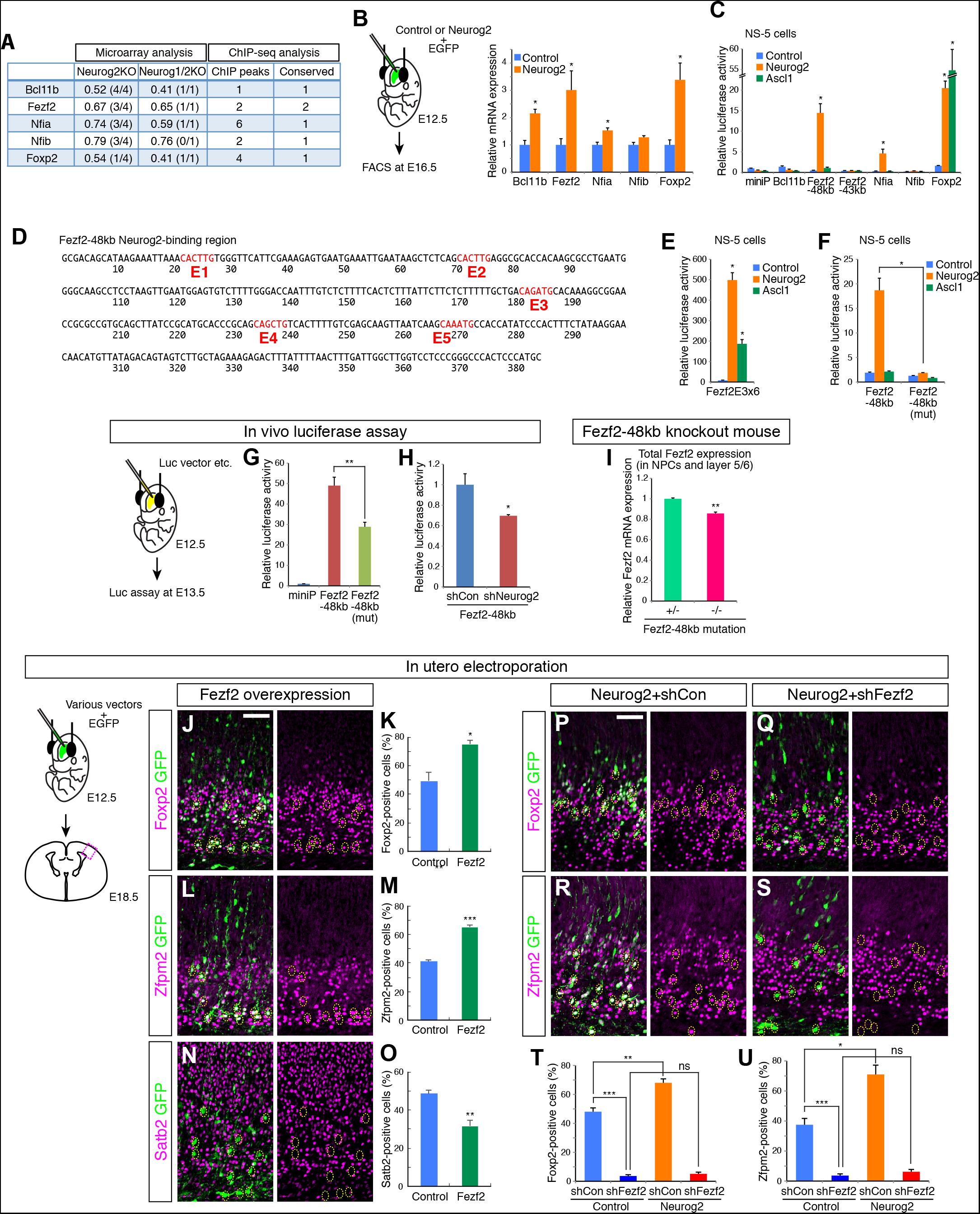
*Fezf2* is a target of *Neurog2* for cortico-thalamic neuron specification. (A) A list of candidate *Neurog1/2* target genes, that have decreased expression levels in *Neurog2* and *Neurog1/2* mutant cortices and Neurog2 ChIP-Seq peaks in their vicinity. The numbers of conserved ChIP-Seq peaks in each gene between mouse and human are also shown. (B) The mRNA levels of the indicated genes, normalized by that of *Ppia,* are shown. Samples were collected by FACS from E16.5 cortices that had been electroporated at E12.5 with *EGFP* or *Neurog2* expression vectors (Control, n = 3; *Neurog2,* n = 3). (C) Trancriptional activation of Neurog2-binding genomic elements by *Neurog2* and *Ascl1.* Neurog2-binding elements were cloned into a luciferase vector harboring a minimal promoter and transfected in NS5 cells together with control, *Neurog2* or *Ascl1* expression vector, and their luciferase activities were measured. Luciferase activity was normalized by the *Renilla* luciferase activity (Control, n = 3; *Neurog2,* n = 3; *Ascl1* n = 3). (D) The DNA sequence of the Neurog2-binding region 48kb upstream of the transcription start site of the *Fezf2* gene contains five E-box motif (E1–E5). (E) The transcriptional activity of tandem repeats of the E3 motif was measured as in (C) (Control, n = 3; *Neurog2,* n = 3; *Ascl1* n = 3). (F) The activity of the *Fezf2–48kb* element with mutations in the E3 motif [*Fezf2–48kb(mut)*] was measured as in (C) (n = 3 in each condition). (G) E12.5 cortices were electroporated with the indicated luciferase vectors. After 1 day, the brains were lysed and subjected to luciferase analysis (miniP, n = 3; *Fezf2–48kb,* n = 4; *Fezf2–48kb(mut),* n = 4). (H) E12.5 cortices were electroporated with the *Fezf2–48kb* luciferase vector together with a control shRNA (*shCon*) or *Neurog2* shRNA (*shNeurog2*) (*shCon,* n = 4; *shNeurog2,* n = 4). (I) Expression level of *Fezf2* mRNA in E14.5 cortices of control or *Fezf2–48kb* element knockout mice (heterozygous control, n = 3; homozygous mutant, n = 5). (J–O) Immunostaining for the indicated subtype-specific markers in E18.5 cortices after *in utero* electroporation at E12.5 of *EGFP* and *Fezf2* expression vector (J, L, and N). Quantification of the percentage of marker-positive cells among EGFP-positive cells (K, M, and O) (for Foxp2 and Satb2, Control, n = 4; *Fezf2,* n = 3 | for Zfpm2, Control, n = 4; *Fezf2,* n = 4). (P–U) Experiments were performed as in (J)–(M) with transfection of the indicated plasmids (*shCon* + Control, n = 4; *shFezf2* + Control, n = 4; *shCon* + *Neurog2,* n = 3; *shFezf2* + *Neurog2,* n = 4). Scale bars: 50 μm (J and P). Data are represented as means ± SEM; \**p* < 0.05; *\*\*p* < 0.005; *\*\*\*p* < 0.0005 from two-tailed unpaired Student’s *t*-test with Welch’s correction; ns, not significant.

To provide further evidence for direct regulation by *Neurog2,* we determined whether the Neurog2 binding sites identified by ChIP-Seq actually mediated *Neurog2*-dependent transcription. We cloned in a luciferase reporter construct Neurog2-bound regions that are well conserved between mouse and human (ECR browser (Ovcharenko et al., 2004)), since important regulatory elements such as enhancers tend to be conserved between mammalian species (Elnitski et al., 2003; Loots et al., 2000). The conserved, Neurog2-bound genomic regions associated with *Fezf2* [48kb upstream of *Fezf2* transcription start site (TSS)] and *Nfia* (345kb upstream of *Nfia* TSS) induced luciferase activity upon *Neurog2* expression in the neural stem cell line NS-5 (Figure 3C). In contrast, Ascl1, another proneural bHLH transcription factor, did not promote transcription in these assays, while both *Neurog2* and *Ascl1* activated transcription when interacting with an element in the *Foxp2* gene (0.8kb upstream of the *Foxp2* TSS) (Figure 3C).

The transcription factor Fezf2 has previously been implicated in the specification of CTNs in layer 6 (Chen et al., 2005a; Molyneaux et al., 2005) and of subcerebral projection neurons in layer 5 (Chen et al., 2005a; Chen et al., 2005b; Molyneaux et al., 2005). Since *Fezf2* expression was reduced in *Neurog2* and *Neurog1/2* mutant cortices (Figure 3A) and was induced by overexpression of *Neurog2* (Figure 3B), and since Neurog2 was bound to a conserved element 48kb upstream of Fezf2 TSS (Figure 3A) and could specifically induce transcription when interacting with this element (Figure 3C), *Fezf2* appeared to be a good candidate to mediate some of the activity of *Neurog* genes in specification of CTNs. We found five consensus binding motifs for bHLH factors (E-boxes) in the *Fezf2–48kb* genomic sequence that responded to *Neurog2* expression in the luciferase assay (Figure 3D). The third motif (E3) matches perfectly the E box CA**<u>GA</u>**TG which is specifically recognized by Neurogenin proteins (Seo et al., 2007) and is present in *Neurogenin-regulated* enhancers in the *Neurod1* and *Rnd2* genes (Heng et al., 2008; Huang et al., 2000). A luciferase construct harboring tandem repeats of the E3 motif was strongly activated by *Neurog2* (Figure 3E) and point mutations disrupting this E-box abolished the activation of the *Fezf2–48kb* element by *Neurog2* (Figure 3F), thus providing strong evidence that E3 mediates most of the *Neurog2*-dependent activity of this enhancer.

We next assessed the activity of the *Fezf2–48kb* enhancer in the embryonic cortex and determined the role of *Neurog2* in activating this element *in vivo.* IUE of the *Fezf2–48kb* luciferase reporter construct in cortical NPCs at E12.5 strongly induced luciferase activity *in vivo* 1 day later, when most of the electroporated cells were still located in the VZ/SVZ (Figure 3G). The mutation of the E3 motif reduced the activity of the *Fezf2–48kb* element by 41% (Figure 3G), and *Neurog2* knockdown by co-electroporation of a *Neurog2* shRNA construct decreased the activity of the wild-type element by 30% (Figure 3H), demonstrating that endogenous Neurogenin factors contribute to the activity of this enhancer *in vivo.*

To further investigate whether *Fezf2–48kb* regulates the *Fezf2* gene in the embryonic cortex, we generated a mouse line in which this putative enhancer has been disrupted (*Fezf248kbenhKO* mutation) using the CRISPR/Cas9 system (Cong et al., 2013; Mali et al., 2013; Yang et al., 2013) (see Methods). The mRNA level of *Fezf2* in whole cortices of E14.5 mice homozygous for the *Fezf248kbenhKO* mutation was significantly reduced compared to that in heterozygous controls (Figure 3I). *Fezf2* is expressed at much higher levels in subcerebral layer 5 projection neurons than in CTNs (http://developingmouse.brain-map.org and data not shown) and has been reported to be controlled by multiple enhancers (Han et al., 2011; Kwan et al., 2008; McKenna et al., 2011; Shim et al., 2012). The limited reduction of *Fezf2* expression in *Fezf248kbenhKO* mice could thus be due to its remaining expression in layer 5 subcerebral projection neurons. Collectively, these results indicate that Neurog1/2 proteins control *Fezf2* expression in CTNs at least in part by interacting with its −48kb enhancer.

### *Neurog2* induces *Fezf2* to activate the cortico-thalamic neuron specification programme

We next examined whether *Fezf2* acts downstream of *Neurogenins* for the specification of CTNs. For this, we first showed that *Fezf2* overexpression by IUE promotes CTN specification in the same way as *Neurog2* does. Electroporation of *Fezf2* at E12.5 increased the generation of Foxp2-, Zfpm2-, Bcl11b- and Nfia-expressing CTNs (Figures 3J–3M; Figures S3A–S3D) and decreased the generation of Satb2-positive CCNs (Figures 3N and 3O).

To determine whether *Fezf2* mediates the specification of CTNs by *Neurog2,* we examined the consequence of suppressing *Fezf2* expression in Neurog2-overexpressing cells. We used a *Fezf2* knockdown construct (Chen et al., 2005b) that efficiently suppresses *Fezf2* expression (Figure S3E) and prevents the generation of CTNs when electroporated in E12.5 embryonic cortices, as expected from the known role of *Fezf2* in CTN specification (Chen et al., 2005a; Molyneaux et al., 2005) (control experiments in Figures 3T and 3U). Co-electroporation of *Neurog2* with the *Fezf2* knockdown construct failed to induce the generation of CTNs, in contrast to electroporation of *Neurog2* alone (Figures 3P–3U; Figures S3F and S3G). Therefore, *Fezf2* acts downstream of the *Neurogenin* genes and is essential for their activity of specification of the CTN fate.

### *Neurog2* overexpression fails to induce cortico-thalamic neurons after E12.5

The finding that *Neurog1/2* specify the subtype identity of CTNs enabled us to study the mechanisms controlling the temporal specification of this early-born population of cortical neurons. In particular we asked the question of why the *Neurogenin-induced* programme of CTN specification is terminated when NPCs stop generating layer 6 neurons after E12.5, although *Neurog1/2* remain expressed throughout cortical neurogenesis. First, we asked whether overexpression of *Neurog2* at E13.5 overrides the normal specification of E13.5 NPCs to layer 5 and 4 fates and reprogrammes these cells to a CTN fate. IUE of *Neurog2* into E13.5 cortices did not induce CTN markers (Figures 4A–4D, 4G, and 4H). Instead, most *Neurog2-*expressing cells, like control EGFP-electroporated cells, expressed the CCN marker Satb2 (Figures 4E, 4F, and 4I). This result suggests that a mechanism restricting the *Neurog1/2-*induced CTN specification programme is already in place in E13.5 NPCs.

**Figure 4.**
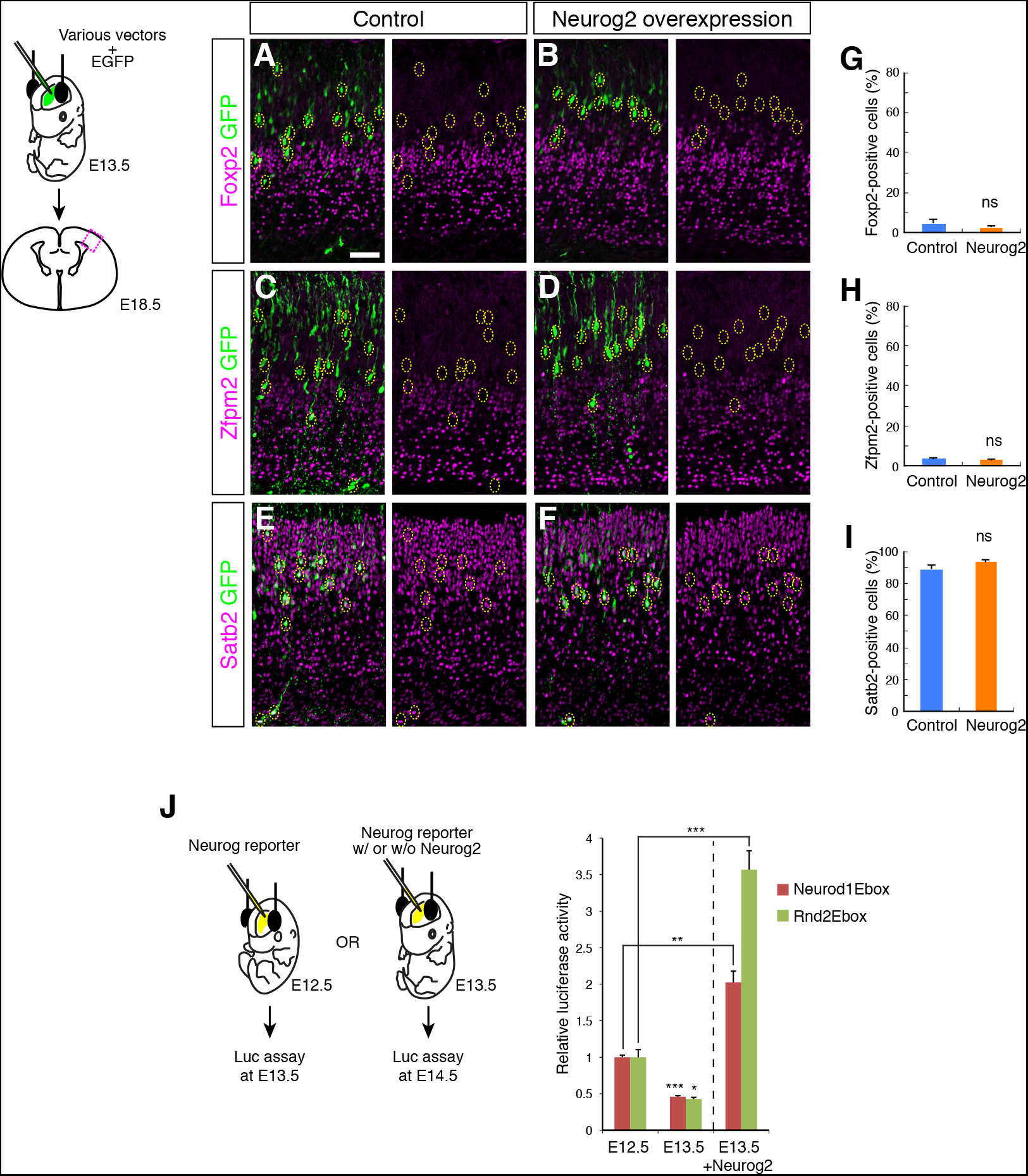
*Neurog2* overexpression in E13.5 NPCs fails to induce cortico-thalamic neurons. (A–F) Immunostaining for the indicated subtype-specific markers in E18.5 cortices after *in utero* electroporation at E13.5 of *EGFP* and control (A, C, and E) or *Neurog2* expression vector (B, D, and F). (G–I) Quantification of the percentage of marker-positive cells among EGFP-positive cells in (A)–(F) (Control, n = 4; *Neurog2,* n = 4). (J) E12.5 or E13.5 cortices were electroporated with *Neurog* reporters containing tandem repeats of E-box motifs (Neurod1Ebox, Rnd2Ebox). E13.5 cortices were also co-electroporated with *Neurog2* expression vector together with the *Neurog* reporters. After 1 day, the brains were lysed and subjected to luciferase analysis (for Neurod1Ebox: E12.5, n = 5; E13.5, n = 6; E13.5 with Neurog2, n = 4; for Rnd2Ebox: E12.5, n = 4; E13.5, n = 4; E13.5 with Neurog2, n = 4). Scale bar: 50 μm. Data are represented as means ± SEM; *\*p* < 0.05; \*\**p* < 0.005; *\*\*\*p* < 0.0001 from two-tailed unpaired Student’s t-test with Welch’s correction; ns, not significant.

An alternative explanation of the above data could be that *Neurog2* is inactive in E13.5 NPCs. A previous study has indeed shown that the neuronal differentiation-promoting activity of *Neurog2* is significantly lower in E14.5 NPCs than E12.5 NPCs, due to phosphorylation by the GSK3 kinase that interferes with the binding of Neurog2 to its targets (Li et al., 2012). To determine whether this mechanism interferes with the CTN specification function of *Neurogenins,* we measured endogenous *Neurogenin* activity in E12.5 and E13.5 cortices by electroporating reporter constructs consisting of concatemerised E-boxes from the *Neurod1* and *Rnd2* genes, two targets of *Neurog2,* driving the luciferase reporter. There was a significant reduction of the activity of the E-box reporters between E12.5 and E13.5 (Figure 4J), suggesting that the transcriptional activity of Neurog1/2 indeed declines immediately after the period of generation of CTNs. ChIP-PCR analysis of *Neurog2* targets also showed that p300 recruitment, but not Neurog2 binding, became weaker between E12.5 and E14.5, suggesting that Neurog1/2 have a reduced capacity to recruit co-activators after the period of CTN production (Figure S4). However, when *Neurog2* was co-electroporated with E-box reporters at E13.5, *Neurogenin* activity was increased to a level above endogenous *Neurogenin* activity at E12.5, which is sufficient to induce CTN fate during normal development (Figure 4J). Therefore, the inability of overexpressed *Neurog2* to induce CTNs at E13.5 is not due to a reduction of its transcriptional activity. This in turn suggests that a mechanism other than the attenuation of endogenous Neurogenin activity blocks the induction of the CTN programme by *Neurog1/2* at this stage.

### PRC2 components are required to restrict the cortico-thalamic neuron fate to layer 6 neurons

The fact that *Neurog2* overexpression at E13.5 strongly activated E-box reporters (Figure 4J) but failed to induce CTNs (Figures 4A–4I) suggested that the fate restriction of E13.5 NPCs might involve epigenetic rather than transcriptional mechanisms. Recent studies have revealed a contribution of components of Polycomb Repressive Complexes (PRCs) to the timing of generation of different cell types in the neocortex, although whether PRC proteins play a role in the temporal restriction of the CTN specification programme has not been addressed (Morimoto-Suzki et al., 2014; Pereira et al., 2010).

To determine whether PRCs control the timing of generation of CTNs, we first targeted *Eed,* an essential component of PRC2, using efficient shRNAs (Cooper and Brockdorff, 2013) (Figure S5A). While overexpressing *Neurog2* alone at E13.5 did not induce CTN markers, simultaneously overexpressing *Neurog2* and silencing *Eed* by IUE of *Neurog2* and either of two different *shEed* constructs resulted in a marked increase in expression of CTN markers (Figures 5B, 5C, 5E, 5F, 5J, and 5K; Figures S5C–S5G). Moreover, the expression of the CCN marker Satb2 was decreased by this treatment (Figures 5H, 5I, and 5L) and many axons from *Neurog2* and *shEed* co-electroporated neurons were found in the dorsal thalamus, while only very few axons from neurons electroporated with control vectors were found at this location (Figures 5M–5Q). Therefore, in the absence of *Eed, Neurog2* could reprogramme E13.5 NPCs into proper CTNs harboring appropriate molecular markers and axonal projection patterns.

**Figure 5.**
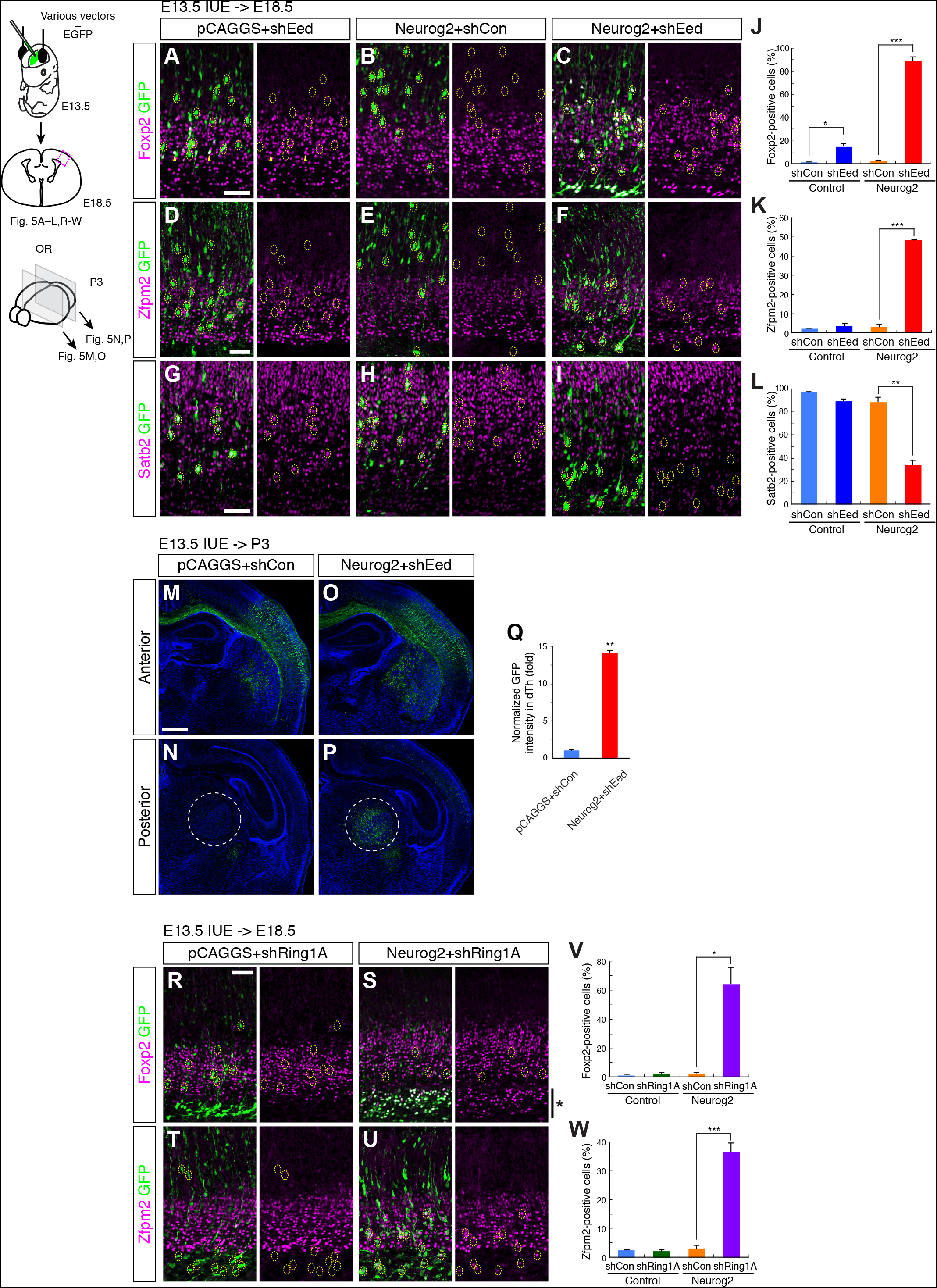
*Neurog2* reprogrammes E13.5 NPCs to a cortico-thalamic fate when PRC2 or PRC1 is inactivated. (A–I) Immunostaining for the indicated subtype-specific markers in E18.5 cortices after *in utero* electroporation at E13.5 of *EGFP* and the indicated plasmids. (J–L) Quantification of the percentage of marker-positive cells among EGFP-positive cells in (A)–(I) (for Foxp2, *shCon* + Control, n = 4; *shEed* + Control, n = 4; *shCon* + *Neurog2,* n = 4; *shEed* + *Neurog2,* n = 4 | for Zfpm2, *shCon* + Control, n = 4; *shEed* + Control, n = 4; *shCon* + *Neurog2,* n = 4; *shEed* + *Neurog2,* n = 3 | for Satb2, *shCon* + Control, n = 4; *shEed* + Control, n = 4; *shCon* + *Neurog2,* n = 3; *shEed* + *Neurog2,* n = 4). (M–Q) Immunostaining for EGFP in P3 brains after *in utero* electroporation at E13.5 of *EGFP* and the indicated plasmids. A representative image is shown in each condition. Dotted circles at a posterior level indicate the dorsal thalamus (N and P). Quantification of (M-P) is shown (Q) (Control + *shCon,* n = 7; *Neurog2* + *shEed,* n = 5). (R–W) Experiments were performed as in (A) and (D) with transfection of the indicated plasmids. The asterisk in (S) shows mis-located cells that are positive for Foxp2 (for Foxp2, *shCon* + Control, n = 4; *shRing1A* + Control, n = 4; *shCon* + *Neurog2,* n = 4; *shRing1A* + *Neurog2,* n = 4 | for Zfpm2, *shCon* + Control, n = 4; *shRing1A* + Control, n = 3; *shCon* + *Neurog2,* n = 4; *shRing1A* + *Neurog2,* n = 3). Scale bars: 50 μm (A, D, G, and R); 500 μm (M). Data are represented as means ± SEM; *\*p* < 0.05; \*\**p* < 0.001; *\*\*\*p* < 0.0001 from two-tailed unpaired Student’s *t*-test with Welch’s correction.

To confirm that this phenotype resulted from a loss of PRC2 activity, we also silenced *Suz12* (He et al., 2012), another essential component of PRC2, while overexpressing *Neurog2,* and obtained similar results (Figures S5H–S5N). Collectively, these results indicate that when PRC2 function has been abolished, increased *Neurog2* activity can reprogramme E13.5 NPCs into a CTN fate, a characteristic normally restricted to E12.5 NPCs.

We also examined if silencing PRC2 components alone influenced the subtype specification of E13.5 NPCs. Although knockdown of either *Eed* or *Suz12* increased moderately the expression of Foxp2 (Figures 5A and 5J; Figures S5I and S5M), it did not alter the expression of other CTN markers including Zfpm2 (Figures 5D and 5K; Figures S5K and S5N) and Bcl11b (Figures S5B and S5E) or that of the CCN maker Satb2 (Figures 5G and 5L), suggesting that the loss of PRC2 function alone is not sufficient to differentiate E13.5 NPCs into CTNs.

### Both PRC2 and PRC1 suppress the cortico-thalamic fate in layer 4/5 NPCs

Because of increasing evidence that PRC2 does not always act by recruiting PRC1 and promoting histone modifications (Biggar and Li, 2015), we asked whether PRC1 is also involved in the restriction of the CTN fate. Similar to the silencing of PRC2 components, silencing the PRC1 catalytic subunit *Ring1A* (Jacob et al., 2011) (Figure S5O) and simultaneously overexpressing *Neurog2* in E13.5 NPCs increased expression of CTN markers (Figures 5S and 5U–5W), while it did not have this effect when *Neurog2* was not overexpressed (Figures 5R, 5T, 5V, and 5W). Knockdown of the other PRC1 catalytic subunit, *Ring1B* (He et al., 2008) (Figure S5P) had a similar effect (Figures S5Q–S5V). These results indicate that the restriction of the CTN fate to E12.5 NPCs involves the canonical PRC2-PRC1 pathway catalysing histone modifications.

### The *Foxp2* promoter is temporally regulated by the PRC machinery

What is the mechanism by which PRCs suppress the CTN-inducing activity of *Neurog2* in E13.5 NPCs? We first asked whether PRCs repress the *Fezf2* locus, a direct and essential target of *Neurog2* for CTN specification (Figure 3), which has been reported to be progressively repressed by the PRC1 component *Ring1B* after E14.5 (Morimoto-Suzki et al., 2014). Overexpression of *Neurog2* could up-regulate *Fezf2* expression as efficiently at E13.5 than at E12.5 (compare Figures 6A and 3B), suggesting that the promoter of *Fezf2* is still accessible at E13.5 and that repression of *Fezf2* is unlikely to account for the suppression of the CTN fate at this stage.

**Figure 6.**
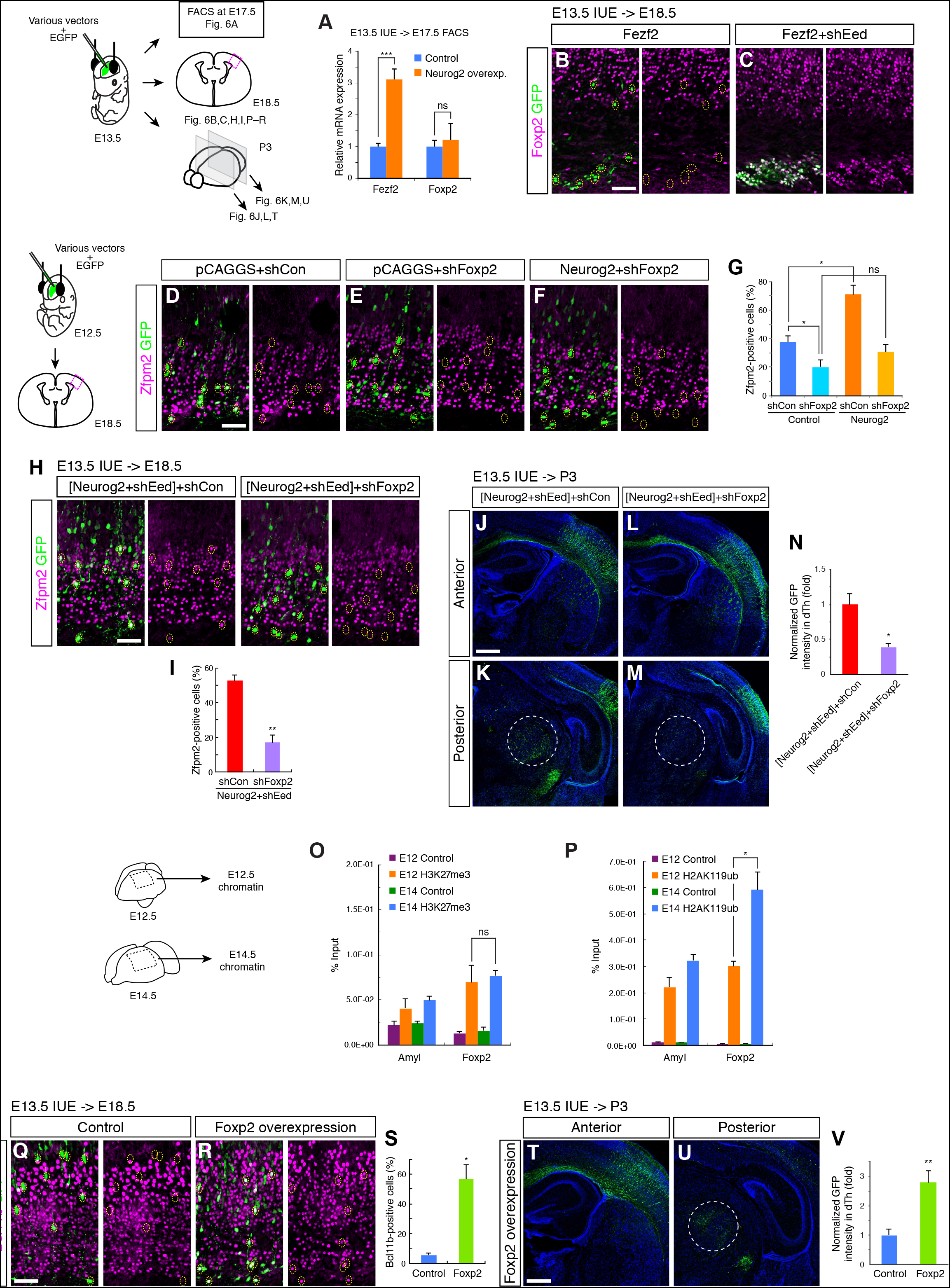
Involvement of *Foxp2* in cortico-thalamic neuron specification and temporal regulation of its promoter by the PRC machinery. (A) mRNA levels of *Fezf2* and *Foxp2.* Samples were collected by FACS from E17.5 cortices that had been electroporated at E13.5 with *EGFP* and control or *Neurog2* expression vector (Control, n = 6; *Neurog2,* n = 4). (B and C) Immunostaining for Foxp2 in E18.5 cortices after *in utero* electroporation at E13.5 of *EGFP* and *Fezf2* with or without *shEed.* (D–G) Immunostaining for Zfpm2 in E18.5 cortices after *in utero* electroporation at E12.5 of EGFP and the indicated plasmids. Quantification is shown in (G) *(shCon* + Control, n = 4; *shFoxp2* + Control, n = 4; *shCon* + *Neurog2,* n = 3; *shFoxp2* + *Neurog2,* n = 4). (H and I) Immunostaining for Zfpm2 in E18.5 cortices after *in utero* electroporation at E13.5 of *EGFP* and the indicated plasmids. Quantification is shown in (I) ([*Neurog2* + *shEed*] + *shCon,* n = 4; [*Neurog2* + *shEed*] + *shFoxp2,* n = 4). (J–N) Immunostaining for EGFP in P3 brains after *in utero* electroporation at E13.5 of *EGFP* and the indicated plasmids. A representative image is shown in each condition. Dotted circles at a posterior level indicate the dorsal thalamus (K and M). Quantification of (J–M) is shown (N) ([*Neurog2* + *shEed*] + *shCon,* n = 4; [*Neurog2* + *shEed*] + *shFoxp2,* n = 4). (O and P) ChIP-qPCR analysis with H3K27me3 antibody (O) or H2AK119ub antibody (P) using chromatin prepared from E12.5 and E14.5 cortices. The amount of promoter regions of *Amyl* (as a control) and *Foxp2* genes in the immunoprecipitated chromatin was detected by qPCR. The percentages of input chromatin were quantified in duplicate from three independent chromatin samples of different litters. (Q–S) Immunostaining for Bcl11b in E18.5 cortices after *in utero* electroporation at E13.5 of *EGFP* and control or *Foxp2* expression vector. The percentage of Bcl11b-positive cells among EGFP-positive cells was quantified (S) (Control, n = 4; *Foxp2,* n = 3). (T–V) Axon trajectories in P3 brains from the neurons that had been electroporated with *EGFP* and *Foxp2* vector at E13.5. A representative image is shown. Dotted circles at a posterior level indicate the dorsal thalamus. Quantification is shown (V) (Control, n = 7; *Foxp2,* n = 4). Scale bars: 50 μm (B, D, H, and Q); 500 μm (J and T). Data are represented as means ± SEM; \**p* < 0.05; \*\**p* < 0.005; \*\*\**p* < 0.0001 from two-tailed unpaired Student’s *t*-test with Welch’s correction.

We next investigated the possibility that step(s) downstream of the induction of *Fezf2* by *Neurog2* in the CTN specification programme might be suppressed by PRCs at E13.5. In support of this hypothesis, overexpressing *Fezf2* at E13.5 did not efficiently induce the CTN fate after 5 days (Figure 6B; Figure S6A), while inactivating PRCs by *Eed* knockdown while overexpressing *Fezf2* markedly increased the expression of CTN markers (Figure 6C; Figure S6B), suggesting that PRCs suppress *Fezf2* activity by targeting its downstream effectors.

Fezf2 is required for expression of *Foxp2* in CTNs (Chen et al., 2005a; Molyneaux et al., 2005) and it binds to the promoter of the *Foxp2* gene (Lodato et al., 2014), suggesting that *Foxp2* is one of *Fezf2* direct transcriptional targets. When *Neurog2* was overexpressed and *Eed* silenced at E13.5, Foxp2 was up-regulated to a level higher than that seen in adjacent non-electroporated neurons (Figures 5A–5C). Moreover, silencing of *Eed* or *Suz12* alone was sufficient to induce Foxp2 expression (Figures 5A and 5J; Figures S5I and S5M), while other CTN markers were not affected. Together, these observations suggested that *Foxp2* might be targeted by PRCs to suppress the CTN fate downstream of *Neurog2* and *Fezf2.*

To address this possibility, we first determined whether *Foxp2* is indeed involved in the specification of the CTN fate downstream of *Neurog2.* Silencing *Foxp2* (Tsui et al., 2013) (Figures S6C and S6D) at E12.5 moderately but significantly reduced the normal generation of CTNs marked by Zfpm2 (Figures 6D, 6E, and 6G) and by axons growing into the dorsal thalamus (Figures S6E–S6I). In addition, *Foxp2* knockdown efficiently blocked the induction of supernumerary CTNs by *Neurog2* overexpression at E12.5 (Figures 6D, 6F, and 6G, and Figure 3R for *Neurog2* + *shCon).* Moreover, knockdown of *Foxp2* in cells overexpressing *Neurog2* and silenced for *Eed* at E13.5 suppressed the reprogramming of electroporated neurons to a CTN fate (Figures 6H and 6I) and the re-direction of axons toward the dorsal thalamus (Figures 6J–6N). Therefore, *Foxp2* is an important determinant of CTN fate downstream of *Neurog2,* during both normal development and reprogramming of layer 4/5 neurons.

Next, we examined whether PRCs directly target *Foxp2* to suppress the CTN fate after layer 6 generation. For this, we investigated PRC-dependent histone modifications at the *Foxp2* promoter both during and after the period of CTN generation (E12.5 and E14.5) using chromatin immunoprecipitation assays. H3K27me3, which is deposited by PRC2, was similarly enriched at the *Foxp2* promoter at E12.5 and E14.5 (Figure 6O). On the other hand, H2AK119ub, which is deposited by PRC1, was significantly more abundant at the *Foxp2* promoter at E14.5 than E12.5 (Figure 6P). This suggests that PRC1 is active at the *Foxp2* promoter and that this activity increases after layer 6 generation.

We next determined whether repression of *Foxp2* transcription by PRCs contributes to the suppression of the CTN fate after E12.5, by asking whether forced *Foxp2* expression could overcome this temporal restriction. Overexpression of *Foxp2* at E13.5 was not able to induce most CTN markers, including Zfpm2 and Nfia (Figures S6J and S6K), but it could strongly induce expression of Bcl11b (Figures 6Q–6S). Moreover, *Foxp2*-expressing neurons extended their axons into the dorsal thalamus (Figures 6T–6V). This suggests that repression of *Foxp2* by PRCs is an important mechanism that suppresses parts of the *Neurog1/2*-induced CTN specification programme after layer 6 neurons have been produced.

## Discussion

In this study, we have examined the mechanisms of temporal specification of mammalian NPCs by focusing on a novel programme of cortico-thalamic neuron specification involving a *Neurog1/2–>Fezf2–>Foxp2* regulatory cascade. We have identified two independent mechanisms that restrict this CTN specification programme to cortical NPCs generating layer 6 neurons, namely the regulation of Neurog1/2 transcriptional activity and an engagement of Polycomb repressive complexes repressing downstream genes in the CTN-specification programme. In E12.5 NPCs, Neurog1/2, which have high transcriptional activity, switch on a *Fezf2*-dependent genetic pathway inducing CTNs. After E12.5, the activity of Neurog1/2 is progressively reduced and simultaneously, some of the downstream genes involved in CTN specification, including *Foxp2*, become inaccessible because of repressive modifications catalysed by PRCs (Figure 7). Each of these mechanisms separately is sufficient to prevent E13.5 NPCs from differentiating into CTNs, revealing a high degree of robustness in the specification of cortical NPCs.

**Figure 7.**
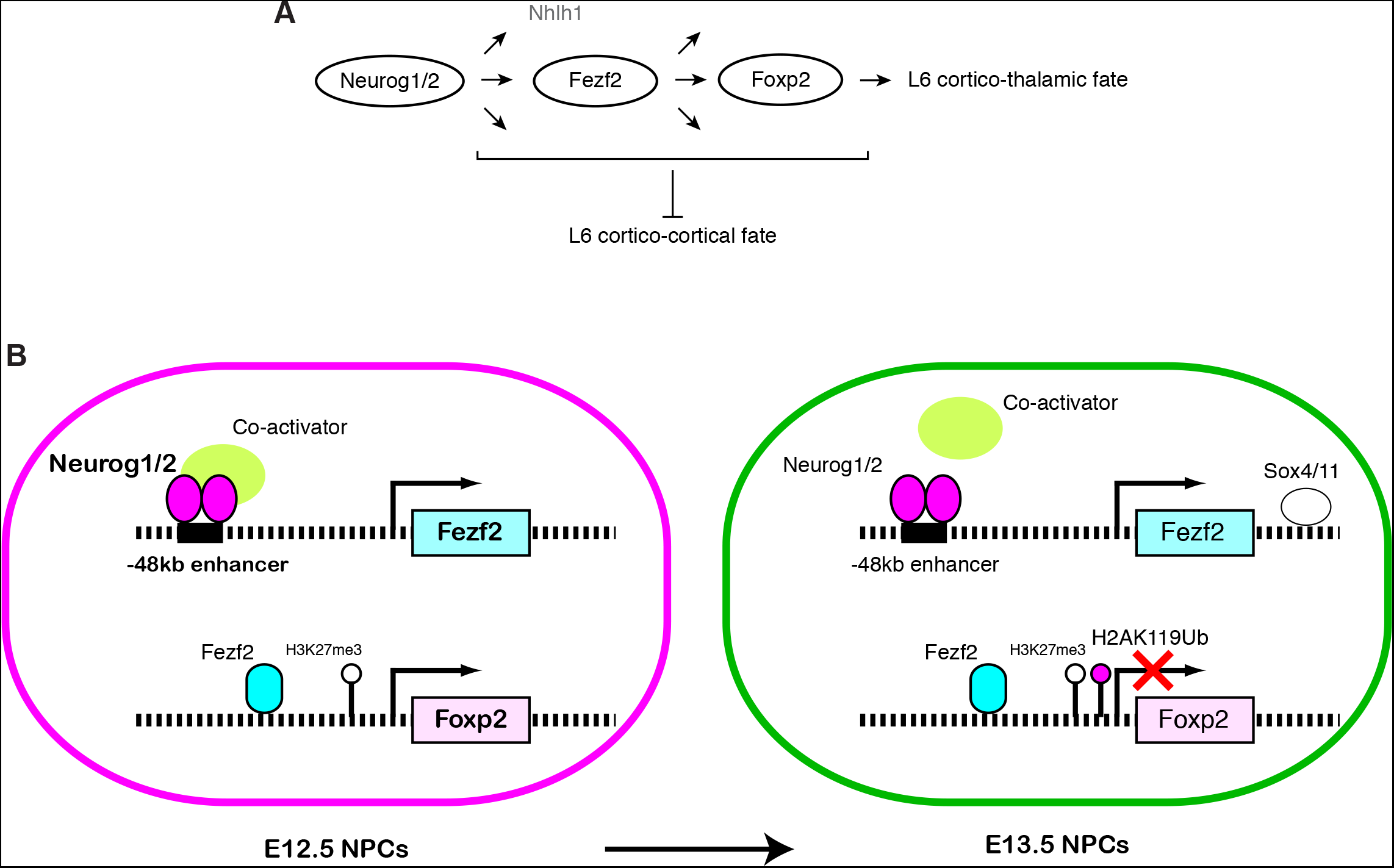
Schematic representation of the regulatory cascade for cortico-thalamic neuron specification and of its temporal control. (A) *Neurog1/2* specify the CTN fate in layer 6 by inducting *Fezf2* expression via direct binding to its enhancer. *Neurog1/2* may also activate other targets including *Nhlh1.* Fezf2, in turn, promotes *Foxp2* expression, presumably by directly activating its promoter. This gene cascade contributes to the establishment of the CTN fate, which antagonizes another major fate of layer 6 neurons, cortico-cortical neurons. (B) In E12.5 NPCs, the gene cascade described in (A) is active. However, this cascade is blocked in NPCs after E12.5 by two independent mechanisms. Neurog1/2 transcriptional activity decreases, presumably because of reduced recruitment of co-activators including p300. CTN fate determinant genes including *Foxp2* become inaccessible because of repressive modifications catalysed by PRC1, which ubiquitinates K119 of Histone H2A.

### Temporal regulation of NPCs in mammalian corticogenesis

Based on extrapolation from studies in *Drosophila,* it has been proposed that the temporal specification of NPCs during mammalian neurogenesis involves the sequential expression of fate determinants (Kohwi and Doe, 2013). However, so far only few transcription factors have been shown to display a temporal pattern of expression during mammalian corticogenesis that is compatible with this model [i.e. Fezf2 (Hirata et al., 2004; Molyneaux et al., 2005), Otx1 (Frantz et al., 1994), Pou3fs (Dominguez et al., 2013) and Ikaros (Alsio et al., 2013)]. Moreover, the functional relevance of theses temporal expression patterns for the subtype specification of NPCs is yet to be determined. In particular, although transplantation experiments have argued for a restriction of the differentiation potential of NPCs during late corticogenesis (Frantz and McConnell, 1996), it is still unclear whether the commitment to a particular neuronal subtype occurs at the NPC or neuronal stage. For example, *Fezf2* is expressed in NPCs at a higher level during early than late cortical development, suggesting a possible function as a temporal determinant of NPC fate. However, *Fezf2* is also expressed in neurons in deep layers and the ectopic expression of *Fezf2* in upper layer neurons is able to reprogramme these neurons to a layer 5/6 subcerebral/subcortical-like fate (De la Rossa et al., 2013; Rouaux and Arlotta, 2013), suggesting that *Fezf2* expression in NPCs may not be necessary for deep layer neuron specification.

Here we demonstrate that *Neurog1/2,* which are expressed mostly by NPCs and only transiently by postmitotic neurons (Hand et al., 2005), control the subtype specification of CTNs, suggesting that this neuronal subtype is specified at the NPC stage. A function of Neurog2 in subtype specification of deep layer cortical neurons has also been recently reported (Dennis et al., 2017). Although *Neurog1/2* are expressed throughout neurogenesis, their subtype specification function operates only in a brief time window during early corticogenesis and must therefore be strictly regulated temporally. Given the lack of evidence for sequentially expressed temporal factors in NPCs, we speculate that target accessibility rather than gene expression might be temporally regulated so that NPCs are specified to different fates at different times.

### *Foxp2* as a temporally regulated target of Polycomb Repressive Complexes

We found in this study that PRCs temporally regulate the *Foxp2* promoter, while in a previous work the PRC1 component *Ring1B* has been shown to temporally regulate the promoter of *Fezf2* (Morimoto-Suzki et al., 2014). *Ring1B* was shown to be required for the termination of subcerebral neuron production at E14.5, concomitant with an increased level of H3K27me3 modification at the *Fezf2* promoter. In contrast to the suppression of the subcerebral fate at E14.5, it seemed unlikely that the suppression of the CTN fate at E13.5 also involves *Fezf2* repression since this gene is active and specifies subcerebral neurons at this stage. Indeed, *Neurog2* could still induce *Fezf2* at E13.5, indicating that the *Fezf2* promoter remains accessible. Instead, we found that the *Foxp2* promoter becomes repressed by PRC1 at E13.5. These results indicate that the repression by PRCs of different targets is differentially controlled temporally, resulting in NPCs becoming restricted to different fates at different stages.

Then, how are PRCs recruited to different genomic loci at different times? We found that PRC1-dependent modifications, but not PRC2-dependent modifications, become more abundant at the *Foxp2* promoter at E14.5 than E12.5, suggesting that PRC1 engagement is temporally regulated. As PRC1 complexes display considerable heterogeneity in composition (Simon and Kingston, 2013), it is possible that a specific composition of the complex enables the targeting of specific loci, in a similar way to what has been proposed for the composition of SWI/SNF or BAF complexes in NPCs versus neurons (Lessard et al., 2007). In addition, PRC1 displacement factors such as Zrf1 (Aloia et al., 2013) could be involved in this target-specific recruitment. Such regulations would make it possible that only particular genes rather than whole specification programmes, are suppressed.

### Temporal regulation of the transcriptional activity of Neurogenins

In addition to the regulation of chromatin accessibility at downstream genes, we found that the reduction of transcriptional activity of Neurog1/2 also contributes to restricting the CTN fate after E12.5, since inactivation of PRC2 or PRC1 could re-specify E13.5 NPCs only when performed together with overexpression of *Neurog2.* It has been recently proposed that a reduction of Neurog2 transcriptional activity by GSK3-mediated phosphorylation accounts for the restriction of *Neurog2* subtype-specification activity to deep layer neurons (Li et al., 2012). However, we found that at E13.5, Neurog1/2 retain significant transcriptional activity but have already completely lost the capacity to induce the CTN fate, indicating that a distinct mechanism operates to suppress the CTN fate. Li *et al.* (Li et al., 2012) also suggested that the reduction of *Neurog2* activity in late corticogenesis is due to an attenuation of Neurog2 binding to its targets. However, we did not observe a significant change in Neurog2 binding between early and late corticogenesis (Figure S4A). Instead, we found reduced p300 recruitment to Neurog2-containing enhancers in late corticogenesis (Figure S4B). Post-translational regulation events [e.g. (Quan et al., 2016)] might also be involved in the progressive weakening of Neurog1/2 activity. Further investigation is required to fully characterise the mechanisms underlying the change in Neurog1/2 activity that occurs as cortical development unfolds.

### Additional targets of Neurog1/2

*Fezf2* has been shown to be necessary and/or sufficient for the expression of several CTN-specific genes, including *Foxp2* and *Zfpm2* (Chen et al., 2005a; Chen et al., 2005b; Molyneaux et al., 2005) but CTNs still retain some of their characteristics in the absence of *Fezf2.* The expression of Bcl11b, a transcription factor expressed broadly by subcortical projection neurons, was not altered by *Fezf2* deletion (Molyneaux et al., 2005; Srinivasan et al., 2012) (see also Figure S3G), and the axons of *Fezf2-deficient* neurons normally reached the dorsal thalamus (Chen et al., 2005a; McKenna et al., 2011). CTNs are therefore less affected by loss of *Fezf2* than by loss of *Neurog1/2,* suggesting that other *Neurogenin* targets might promote *Fezf2-* independent CTN traits. *Nhlh1,* a potential direct target of *Neurog1/2* which is expressed by intermediate progenitors during generation of layer 6 neurons, and is able to induce *Bcl11b* expression when overexpressed at a later stage (K.O. and F.G., unpublished observation), is such a candidate determinant of CTN fate acting downstream of *Neurog1/2* and alongside *Fezf2*

We have shown that *Foxp2* acts downstream of the *Neurog –>Fezf2* pathway in the CTN specification programme and that it is a target of the PRC-mediated repression terminating CTN production. Ectopic *Foxp2* expression was able to induce some CTN traits, including the expression of Bcl11b and a dorsal thalamic axonal trajectory, although it was not sufficient on its own to fully reprogramme late NPCs to a CTN fate. *Fezf2* directly regulates many target genes during cortical development (Lodato et al., 2014) and is likely to induce other components of the CTN specification programme, some of which might also be repressed by PRCs to terminate CTN production.

### Determination of cortico-thalamic or cortico-cortical fate in layer 6

Although CTNs and CCNs are main components in layer 6 (Thomson, 2010; Tasic et al., 2016), their fate choice mechanisms have remained unclear. One possible assumption is a temporal specification mechanism of these population, which was reported recently, although the generation timing of each population is not completely separated but partially overlapping (Hatanaka et al., 2016). Our analysis, in which the fate choice during very short timing was addressed using EdU labelling and electroporation, suggested that the neurons born at around same time actually differentiate either CTNs or CCNs (if temporal specification happens strictly, this only labels CTNs or CCNs). Moreover, Neurog2 overexpression increased CTNs in this short time window, strongly suggesting that Neurogenin activity directly regulates the choice between CTN and CCN fate, but not the timing of differentiation. As Neurog1/2 are expressed by NPCs in an oscillatory manner (Shimojo et al., 2008), we speculate that different levels of Neurogenin activity in NPCs might contribute the genesis of CTNs and CCNs; the NPCs with high Neurogenin activity give rise to CTNs.

### Reprogramming cells by “unlocking” multiple mechanisms

Reprogramming somatic cells to a given cell type (e.g. iPS cells) is inefficient and often requires multiple treatments, suggesting that the specification and maintenance of cell fates are robust processes mobilising multiple redundant mechanisms (Srivastava and DeWitt, 2016). We succeeded in reprogramming layer 4/5 NPCs into layer 6 CTN-producing NPCs by simultaneously enhancing Neurog2 activity and suppressing PRC function. Reprograming strategies can help identify essential developmental mechanisms and may contribute in the future to regenerative medicine.

## Experimental Procedures

### Animals and genome editing

Mice were maintained on a 12-h light/dark cycle with free access to food and water, bred and treated according to the guidelines approved by the Home Office under the Animal (Scientific procedures) Act 1986. Protocols detailing the generation and genotyping of the genetically modified mice used have been described previously for *Neurog1^KIGEP^* (Ma et al., 1998) and *Neurog2^KIGFP^* (Seibt et al., 2003).

To generate *Fezf2–48kb* enhancer null mice, the CRISPR-Cas9 system was used as described previously (Wang et al., 2013). Short guide RNAs (sgRNAs) were identified using the MIT CRISPR Design tool to target the upstream and downstream regions of the *Fezf2–48kb* enhancer, and a pair of oligonucleotides for each targeting site (Table S1) was inserted into px330, a bicistronic expression vector expressing Cas9 and sgRNA. Cas9 mRNA and the sgRNAs were produced and purified as described previously (Yang et al., 2013). Cas9 mRNA (100 ng/ml) and two sgRNAs (25 ng/ml) were injected into the cytoplasm of fertilized eggs. The deletion was confirmed by Sanger sequencing.

### Plasmids

The *Neurog2* expression vector was described previously (Heng et al., 2008). *Fezf2* cDNA (MC202571, Origene) and *Foxp2* cDNA (IMAGE ID 6851153, Source Bioscience) were subcloned into a pCAGGS plasmid. The Neurog2-binding regions were amplified by PCR from the mouse genome and subcloned into pGL4.23 to obtain firefly luciferase reporter constructs. To generate a mutation in the *Fezf2–48kb* E3, the E-box sequence of **<u>C</u>**<u>AGA</u>**<u>TG</u>** was changed into **<u>T</u>**<u>AGA</u>**<u>CT</u>**. The luciferase construct harboring tandem repeats of the E3 motif of *Fezf2–48k* and the *Neurog* reporters harboring *Neurod1* or *Rnd2* E-box were generated by inserting annealed oligonucleotides into pGL4.23 (Table S1). The shRNA constructs including *shFezf2, shEed, shEed#2, shSuz12, shRing1A, shRing1B,* and *shFoxp2* were also generated by inserting annealed oligonucleotides into pSuper-neo (Table S1).

### Immunohistochemistry

Brains removed from embryos and pups were fixed for 1 h in phosphate-buffered saline (PBS) containing 4% PFA (wt/vol), incubated overnight at 4°C with 20% sucrose in PBS (wt/vol), embedded in OCT compound (Sakura Finetek), and sectioned with a cryostat to obtain 14-μm-thick coronal sections.

For primary antibodies, we used mouse antibody to rabbit, mouse and chick antibodies to EGFP (Molecular Probes, A11122; Santa Cruz, sc-9996; Abcam, ab13970), rat antibody to Bcl11b/Ctip2 (Abcam, ab18465), mouse antibody to Satb2 (Abcam, ab51502), rabbit antibody to Tbr1 (Abcam, ab31940), rabbit antibody to Zfpm2/Fog2 (Santa Cruz, sc-10755), rabbit antibody to Nfia (Aviva Systems Biology, OAAB10433), rabbit antibody to Sox5 (Abcam, ab26041), goat antibody to Foxp2 (Santa Cruz, sc-21069), rabbit antibody to Darpp32 (Chemicon, AB1656) and rabbit antibody to Nurr1 (Santa Cruz, sc-990). For detection of Bcl11b, Satb2, Zfpm2, Nfia and Sox5, antigen retrieval was performed by incubating the sections for 20 min at 90°C in 0.01 M sodium citrate buffer (pH 6.0). Because EGFP fluorescence disappeared by antigen retrieval treatment, EGFP was immunostained with chick or moue antibody against EGFP for re-visualization. Immune complexes were detected with Alexa Flour–conjugated secondary antibodies (Molecular Probes). For nuclear staining, 2 μg/ml DAPI (Molecular Probes) was used. Images were acquired using a confocal microscope (SP5, Leica).

### *In utero* electroporation and EdU administration

*In utero* electroporation of the cerebral cortex was performed in anesthetized time-pregnant mice, as previously described (Pacary et al., 2011). Cortices were electroporated with five 50-ms electrical pulses at 31V with 1-s intervals using 3-mm platinum electrodes (Harvard Apparatus). For injection of EdU (5-ethynyl-2’-deoxyuridine), the pregnant mice were injected intraperitoneally with a EdU solution (Santa Cruz, 20 mg per kg body weight). EdU was detected using ClickiT® EdU Imaging kit (Invitrogen).

### RNA isolation and qRT-PCR

Total RNA was extracted using the Nucleobond RNA extraction kit (Macherey Nagel), and subjected to reverse transcription using the High-Capacity RNA-to-cDNA™ kit (Applied Biosystems). Gene expression was detected using TaqMan Gene expression assays (Applied Biosystems) and a 7500 real-time PCR system (Applied Biosystems). Data was analyzed using standard protocols to calculate relative expression with the ddCT method using *Ppia* as an endogenous control. Each probe was analyzed in duplicates for at least 3 independent samples per group.

### Cell culture and transfection

Mouse teratocarcinoma P19 cells were cultured in high-glucose DMEM (Invitrogen) supplemented with 10% fetal bovine serum, 2 mM glutamine and 1% penicillin/streptomycin. Mouse neural stem cell line NS-5 cells were cultured in Euromed-N medium (Euroclone) supplemented with 1% N2 (Invitrogen), epidermal growth factor, basic fibroblast growth factor (Peprotech, both at 10 ng/ml), and 2 μg/ml laminin (Sigma). Cells were plated in 24-well or 48-well plates and transfected with Lipofectamine 2000 reagent according to the manufacturer’s protocol (Invitrogen). Twenty-four hours after transfection, cells were collected for subsequent analyses.

### Luciferase assay

NS-5 cells and cortical cells were transfected with various firefly luciferase constructs together with pGL4.74 (a thymidine kinase promoter-driven *Renilla* luciferase reporter).

The luciferase activities of the cell lysates were measured using the Dual-Luciferase reporter assay system (Promega). The firefly luciferase activity was normalized by the *Renilla* luciferase activity. For *in vivo* luciferase assay, cortical cells were electroporated with various luciferase vectors, the control *Renilla* luciferase vector and *EGFP* expression vector. EGFP-positive areas of the electroporated brains were dissected out and lysed.

### Chromatin immunoprecipitation (ChIP) assay

The dorsolateral portions of the neocortex were dissected out, fixed sequentially with di(N-succinimidyl)-glutarate and 1% formaldehyde in phosphate-buffered saline, lysed in SDS lysis buffer (0.5% SDS, 10 mM EDTA, 50 mM Tris-HCl pH 8.0) supplemented with protease inhibitor cocktail (Roche), sonicated for 20 min. Immunoprecipitations were performed as described previously (Martynoga et al., 2013). Quantities of immunoprecipitated DNA were calculated by comparison with a standard curve generated by serial dilutions of input DNA using a 7500 real-time PCR system (Applied Biosystems) and a SYBR green-based kit for quantitative PCR (iQ Supermix, Bio-Rad). Antibodies and amounts used for 20 μg chromatin: rabbit anti-H3K27me3 (0.25 μg, Millipore, 07–449), rabbit anti-H2AK119ub (0.25 μg, Cell Signaling, D27C4), goat anti-Neurog2 (0.5 μg, Santa Cruz, sc-19233) or rabbit anti-p300 (0.5 μg, Santa Cruz, sc-585). The primers used are summarized in Table S1.

### Quantification of axon projection

The EGFP fluorescence signals in the ventrobasal nucleus of the dorsal thalamus of three or four sections were measured, and the average value was considered as the axon amount in the dorsal thalamus. The values were then normalised by the similarly averaged EGFP fluorescence signals in the somatosensory area of the cortex to adjust the electroporation efficiency among different brains.

### Statistical analysis

Unless indicated otherwise, data were represented as means ± SEM or SD of values from at least three embryos from at least two litters. For quantification of *in vivo* cell counting, all positive cells were counted in the areas where rostro-caudal and medio-lateral levels were carefully matched between animals. For two-group comparisons with equal variance as determined by the *F*-test, an unpaired Student’s *t* test was used. Welch’s correction was used for unpaired t-tests of normally distributed data with unequal variance. Differences between groups were considered to be significant at *p* < 0.05. Each p value was stated in figures or figure legends.

## Acknowledgments

We gratefully acknowledge Sophie Wood and Graham Preece for technical support, Rekha Subramaniams and Jacek Mor for managing the mouse colony, and members of the Guillemot lab for discussions. The study was conceived and the manuscript was written by K.O. and F.G. K.O. performed most of the work while D.L.C.B. helped with ChIP experiments. K.O. was supported by fellowships from the Uehara memorial foundation, the UK Medical Research Council (MRC) and the Francis Crick Institute and D.L.C.B. by fellowships from the Federation of European Biochemical Societies and the Francis Crick Institute. This work was supported by the Francis Crick Institute, which receives its core funding from Cancer Research UK (FC0010089), the UK Medical Research Council (FC0010089), and the Wellcome Trust (FC0010089); by the UK Biotechnology and Biological Sciences Research Council (project grant BB/K005316/1 to F.G.); by the UK Medical Research Council (project grant U117570528 to F.G.); by the MEXT/JSPS (KAKENHI JP17K07061 to K.O.); by the Takeda Science Foundation (project grant to K.O.), and by the Wellcome Trust (investigator award 106187/Z/14/Z to F.G.). The authors declare no conflict of interest.

